# Pressure–distance curves of identical and dissimilar lipid membrane surfaces in water

**DOI:** 10.1101/2025.06.02.656637

**Authors:** Lukas Bange, Olaf Soltwedel, David-Elisa Seibel, Regine von Klitzing, Emanuel Schneck

## Abstract

Investigations of the hydration repulsion between hydrophilic soft interfaces in water, in particular between lipid membranes, rely on accurate experimental measurements of the associated pressure–distance curves. Conventional experimental approaches face challenges especially when it comes to the pressure–distance curves between dissimilar surfaces, a scenario with particular value for the study of the mechanisms responsible for the hydration repulsion. Here, we present an alternative approach based on solid-supported inverse lipid bilayers (ILBs) in which two hydrophilic monolayer surfaces face each other across a thin water layer, as evidenced through x-ray reflectometry. The water uptake as a function of the dehydrating osmotic pressure is precisely measured with the help of ellipsometry under controlled humidity conditions. The measurements reproduce the known hydration decay lengths of interacting phospholipid membrane surfaces and of interacting glycolipid membrane surfaces. In addition, we present pressure–distance curves of the interaction between two dissimilar membrane surfaces, with phospholipids on one side and glycolipids on the other side. These unique measurements of asymmetrical interaction scenarios result in a curve that is very similar to that of two interacting glycolipid membrane surfaces, which can be rationalized on the basis of our current knowledge of the repulsion mechanisms.

## Introduction

Lipid membranes have numerous biological functions.^1^ In the congested physiological environment their functions depend on interactions with other membranes. ^2,3^ While attractive interactions promote membrane adhesion and fusion, repulsive interactions prevent direct membrane contacts and facilitate the release of newly formed vesicles. Range, strength, and sign of the interactions are the net result of a variety of physical mechanisms including dispersion forces, screened electrostatic forces, and undulation forces. ^4,5^ At short separations below 2 - 3 nm, strong repulsion, commonly termed *hydration repulsion*, occurs even for charge-neutral membranes. In a different context this short-range repulsion was discussed already by Langmuir^6^ and it has been known as “steric repulsion” in the colloidal science community. ^7,8^ The hydration repulsion between biological membranes has an essential biological role, as it creates a barrier against close membrane contacts and thereby suppresses uncontrolled membrane adhesion and fusion. ^9^ It was first quantified experimentally for multi-bilayers of charge-neutral phospholipids^10^ and later reproduced with the surface force apparatus (SFA) method.^11^ The hydration repulsion decays approximately exponentially as a function of the membrane separation with typical decay lengths between 0.1 and 0.3 nm^3,10–12^ and has since then been commonly used in modeling the forces between lipid membranes in water.^13,14^ The physical mechanisms underlying the hydration repulsion have been subject to controversial debates since the 1980s. The repulsion has been attributed to the unfavorable overlap of interfacial water layers^15^ and to a decrease in the configurational entropy of the lipid molecules upon surface approach. ^7^ Later studies based on molecular dynamics (MD) simulations have provided additional insights, demonstrating that the characteristics of the hydration repulsion depend on the phase state of the membranes^16^ and on their surface chemistry.^17^ Since, inside cells, membrane surfaces of various lipid compositions frequently get into close proximity, asymmetrical interaction scenarios are highly relevant also from a biological perspective. The interaction between lipid membranes in the aqueous environment is commonly quantified in terms of pressure–distance curves, where the interaction pressure Π is measured as a function of the separation (or water layer thickness) *D*_w_ between the membranes. At prescribed temperature *T* and pressure *p*, the interaction pressure Π is related to the derivative of the Gibbs free energy, *G*, with respect to *D*_w_, at constant chemical potential of water, *μ*(*p,T*)

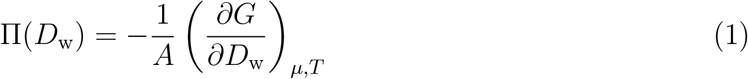

where *A* denotes the surface area.

In experiments, pressure–distance curves Π(*D*_w_) are typically obtained by subjecting multi-bilayers to so-called equivalent pressures of known magnitude.^12,18^ These are realized by controlled competition for the water, i.e., by shifting the water chemical potential to lower values. The water layer thickness *D*_w_ is then deduced from the lamellar periodicity measured by X-ray or neutron diffraction. Equivalent pressures can be exerted by bringing the hydrated membranes into contact with aqueous solutions of hydrophilic polymers. The equivalent pressure then coincides with the osmotic pressure afforded by the polymers. Alternatively, the hydration level can be adjusted via vapor exchange with a water reservoir with lowered chemical potential. The equivalent pressure is then given by the relative humidity *h*_rel_, the temperature *T*, and the volume of a water molecule *v*_*W*_ = 0.03 nm^3^:

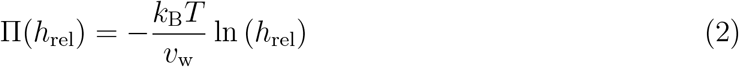

The conventional approaches for measuring pressure–distance curves between lipid layers have some limitations. The multi-bilayer method^10,12^ is very quantitative in that it has a precise definition of the water layer thickness, *D*_w_ = 2*n*_W_*v*_W_*/A*_lip_, where *n*_W_ is the number of water molecules per lipid and *A*_lip_ is the membrane area per lipid.^12,19^ But the method is inapplicable to asymmetrical interaction scenarios because the self-assembly of lipid mixtures does not result in multi-bilayers with surfaces of alternating lipid compositions. The SFA method,^11,20–22^ on the other hand, is applicable to asymmetrical interaction scenarios but does not provide the required precise absolute scale regarding *D*_W_. Instead, assumptions about the lipid layer thicknesses have to be made^11,20,21^ or an adjustable distance offset parameter is employed to interpret the measured pressure–distance curves.

In order to access asymmetrical interaction scenarios with the desired precision, we therefore take an alternative approach in the present work that overcomes the mentioned limitations of the conventional methods: Inverse lipid bilayers (ILBs) with the hydrophilic monolayer surfaces facing towards each other across a thin water layer are deposited onto hydrophobized planar surfaces. Ellipsometry is then used to precisely quantify the water uptake as a function of the relative humidity (i.e., as a function of the equivalent pressure Π). These measurements are fully quantitative and offset-free in *D*_W_ and the ILB architecture can be readily used for asymmetrical monolayer compositions, i.e., for the study of the interaction of chemically dissimilar membrane surfaces. The measurement concept is related to that of a previous study on the hydration of single glycolipid layers ^23^ and to earlier studies on rather long-range repulsive interactions due to either polymer-induced repulsion^24,25^ or electric charges,^25^ which however only covered symmetrical interaction scenarios.

Here, we focus on the hydration repulsion between identical and dissimilar uncharged lipid membrane surfaces. Our measurements reproduce the known pressure–distance curves of interacting identical phospholipid membrane surfaces and of interacting identical glycolipid membrane surfaces, including their pronounced differences. Moreover, we provide the pressure–distance curve of the interaction between two dissimilar membrane surfaces, with phospholipids on one side and glycolipids on the other side. This pressure-distance curve is very similar to that between two glycolipid membrane surfaces, which we rationalize on the basis of the current knowledge of the physical mechanisms responsible for the hydration repulsion. Pressure–distance curves generated with the taken approach are well suited for direct comparison with MD simulations, which can not only predict such curves^16,17,19,26^ but can also be readily applied to both identical and dissimilar membrane surfaces.

## Results and discussion

The inverse lipid bilayers (ILBs) illustrated in Fig. 1 A were prepared by sequentially depositing two lipid monolayers via the Langmuir-Blodgett (LB) technique onto silicon substrates hydrophobically functionalized with OTS (see Methods section for the details). In an architecture that closely mimics the interaction between two lipid membrane surfaces, the two monolayers of the ILB face each other across a thin water layer. Each monolayer is composed of only one type of lipid. Changing the lipid type between the first and the second LB transfer allowed the preparation of asymmetrical interaction scenarios, were monolayers of lipids with different headgroups (HGs) are brought into close proximity.

**Figure 1:**
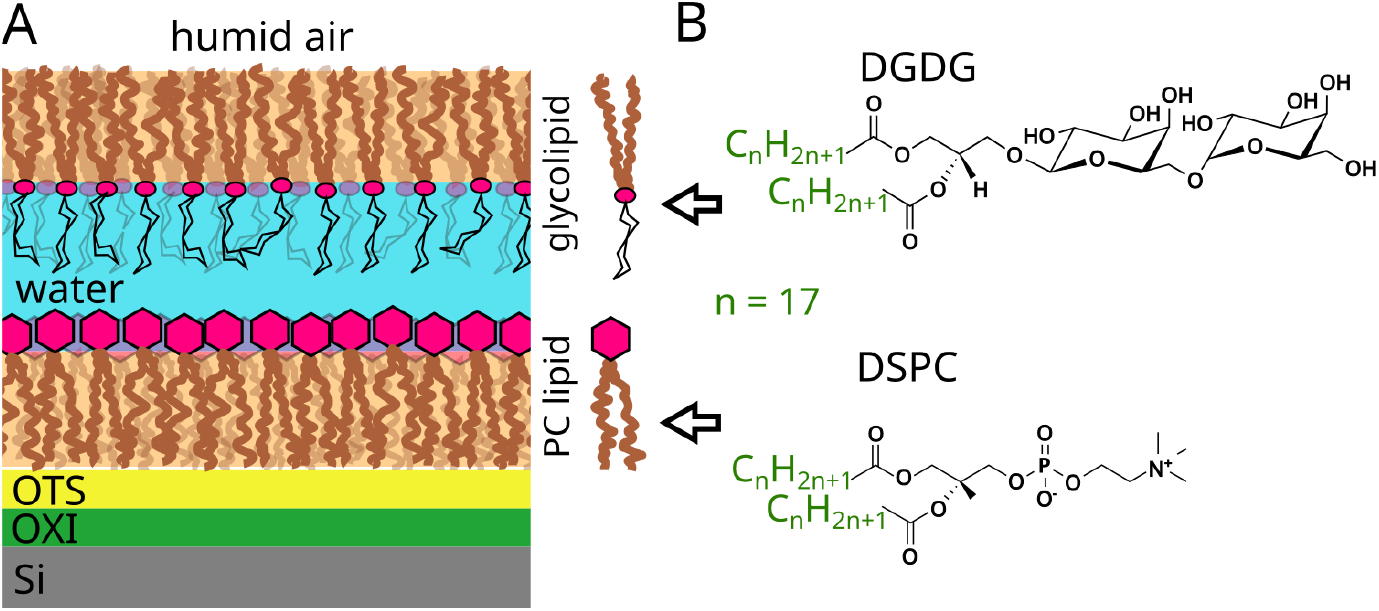
(A) Cartoon of the ILB sample architecture (asymmetrical example with one PC lipid monolayer and one glycolipid monolayer). The ILB is supported by a silicon (Si) surface covered with a layer of thermal oxide (OXI), which is hydrophobically functionalized with OTS. (B) Chemical structures of the phospholipid 1,2-distearoyl-sn-glycero-3-phosphocholine (DSPC) and of the glycolipid digalactosyldiacylglycerol (DGDG).

The chemical structures of the used lipids are shown in Fig. 1 B. Both have saturated C_18_ hydrocarbon chains (HCs) which promote the formation of laterally densely-packed, chain-ordered lipid layers.^27,28^ Lipid exchange between the opposing monolayers in the ILB can be considered negligible because the associated free energy barrier is substantial and lipid mobility is largely suppressed in the chain-ordered state.^2^ The phospholipid DSPC has a zwitterionic phosphatidylcholine (PC) HG, which features a single large single electric dipole. The glycolipid DGDG has a disaccharide HG, which features numerous small electric dipoles in the form of -OH groups. Because of this difference in their HG structure, multi-bilayers of PC lipids retain much more hydration water than multi-bilayers of glycolipids at the same dehydrating osmotic pressure.^3,17^ In other words, PC lipid layers and glycolipid layer exhibit very different pressure–distance curves.

Here, we are primarily interested in realizing equivalent interaction scenarios with our ILB architectures and to determine the pressure–distance curves also in asymmetrical interaction scenarios, where the surface of a layer of PC lipids interacts with the surface of a layer of DGDG. In the following, we will first present an experimental characterization of the ILB architecture by X-ray reflectometry (XRR) and subsequently discuss the pressure–distance curves obtained with ellipsometry at varying osmotic pressures imposed through systematic variations of the relative humidity.

### Sample architecture: XRR

XRR experiments were carried out to determine the interfacial layer structure in terms of its electron density profile. Fig. 2 A shows one of two reflectivity curves measured with a solid-supported ILB composed of DSPC at ambient humidity. This system comprising two PC lipid monolayers was chosen for the XRR measurements because it offers the highest electron density contrast due to the electron-rich P atom in the headgroups. The pronounced intensity oscillations (known also as “Kiessig fringes”) are an immediate proof of the presence of defined layers on the solid surface, and the distance Δ*q*_*z*_ ≈ 0.075 Å^*−*1^ between the first two intensity minima provides a first estimate of the layers’ combined thickness as 2*π/*Δ*q*_*z*_ ≈ 8 nm. The other reflectivity curve, measured with a different sample prepared with the same protocol, is presented in the Supporting Information (Fig. S2) and has virtually identical features, confirming reproducibility.

**Figure 2:**
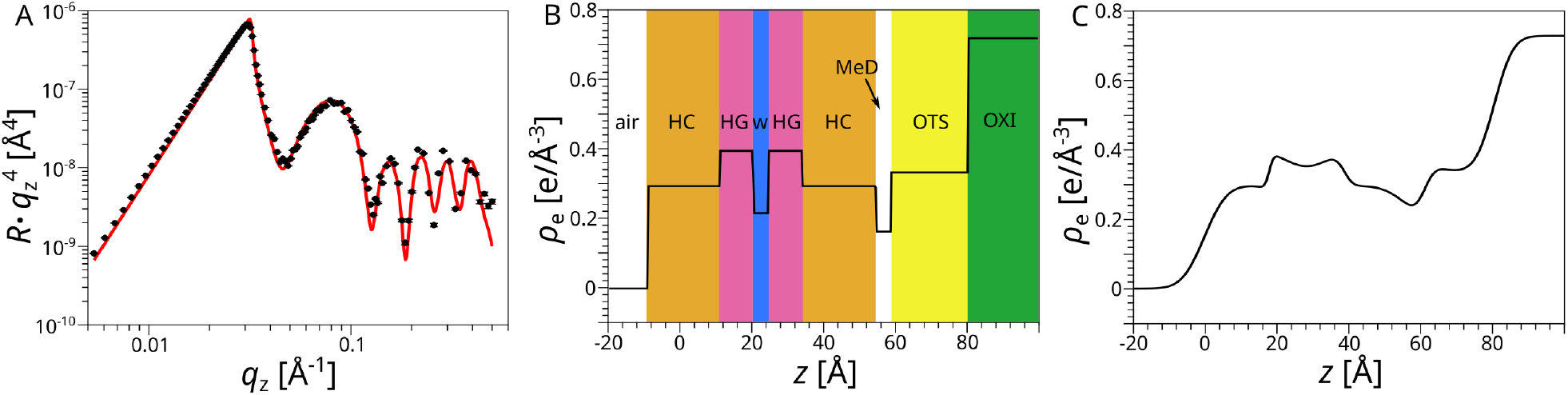
(A) X-Ray reflectivity data from a symmetrical DSPC ILB on an OTS-functionalized silicon chip at ambient humidity. The solid line indicates the simulated reflectivity curve corresponding to the best-matching model parameters. (B) Underlying, roughness free slab model with all relevant layers and their electron densities. HC: hydrocarbon chains; HG: headgroups; W: water; MeD: methyl dip; OXI: thermal silicon oxide. (C) Resulting electron density profile when accounting also for the roughness parameters.

The reflectivity data were analyzed by describing the interfacial electron density profile with a number of homogeneous slabs of adjustable thickness *d* and electron density *ρ*, which represent different portions of the interfacial layers (see Fig. 2 B for a schematic illustration). Starting from the solid Si material including the terminal oxide (OXI), these layers were the OTS layer, a layer of reduced electron density between OTS and the adjacent ILB (termed “methyl dip”, MeD, because this region accommodates the ends of the alkyl chains), and the ILB itself, comprising a proximal (prox, close to the substrate) and a distal (dist, further away from the substrate) monolayer. The ILB was modeled with distinct layers for hydrocarbon chains (HC), headgroups (HG), and an interstitial layer of hydration water (W) between the two headgroup layers. The gradual transitions of the electron density from one layer to the next was described with a set of roughness parameters *σ*. Since *ρ*_OXI_ ≈ *ρ*_Si_,^29^ it was not necessary to include an extra interface between the oxide and the Si bulk material.

Mathematically, the resulting electron density profile can be expressed as

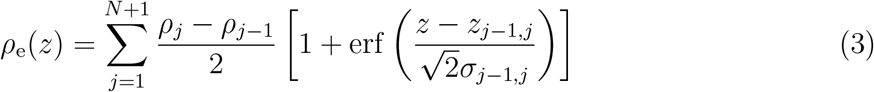

where *ρ*_0_ = 0 (air), *N* is the number of layers, erf(*x*) denotes the error function, *σ*_*j−*1,*j*_ is the roughness of the interface between the (*j*-1)th and the *j*th layer, *z*_0,1_ = 0, and

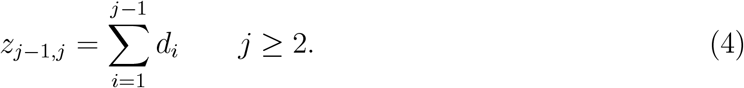

As shown in Fig. 2 B and C, *z* increases when moving from the air towards the silicon support, so that *j* = 0 refers to the bulk medium air and *j* = *N* + 1 to the bulk medium OXI/Si. For reasons of symmetry and in order to reduce the number of free parameters, we set the parameters *d*_HC_, *d*_HG_, *ρ*_HC_, *ρ*_HG_, and *σ*_HC/HG_ of the proximal and distal monolayers to be equal. Moreover, *ρ*_W_ was fixed at the known electron density of pure water in bulk, 0.33 *e*^*−*^*/* Å^3^.

The solid line in Fig. 2 A indicates the simulated reflectivity curve corresponding to the layer-based electron density profile (Eq. 3) with the best-matching parameter set regarding layer thicknesses, electron densities, and roughnesses. The underlying electron density slab model is illustrated in Fig. 2 B, while the associated electron density profile, when also accounting for the roughness parameters, is shown in Fig. 2 C. The simulated reflectivity curve in panel A was calculated from the electron density profile in panel C as described in the Methods section. It can be seen that the experimental reflectivity curve is well reproduced by the model.

The electron density profile in Fig. 2 C clearly exhibits all the expected features, including the two electron-dense headgroup layers separated by a thin hydration layers of lower electron density and the methyl dip between OTS and the HC of the proximal lipid monolayer. The model parameters are summarized in Table 1. With *d*_OTS_ ≈ 2.0 nm, the thickness of the OTS layer is in good agreement with values reported earlier by other groups (≈ 2 - 2.5 nm)^30,31^ and by ourselves (1.7 ± 0.5 nm),^32^ where the variation mainly reflects different definitions of which chemical groups belong to the layer. Good agreement is also found for the thicknesses of the HG layer (*d*_HG_ ≈ 0.9 nm), for which values of 0.7 - 0.8 nm have been reported.^33,34^ Finally, the thickness of the HC layer (*d*_HC_ ≈ 1.8 nm) agrees well with earlier reports (≈ 2 nm)^35^ and with the theoretical prediction *d*_HC_ = *L*_0_ cos *t* ≈ 1.9 nm, where *t* ≈ 30^*°*^ is the chain tilt^28^ and *L*_0_ = 1.54 Å + *N* · 1.26 Å = 22.96 Å is the stretched length of a hydrocarbon chain with *N* = 17 methylene groups,^4,36^ as in DSPC.

**Table 1:** Layer parameters of a DSPC ILB on OTS at ambient humidity as obtained by XRR. MeD: methyl dip; HC: hydrocarbon chains; HG: headgroups; W: water. Error estimates include systematic uncertainties (see Methods section). ^a^Fixed. ^b^*d*_ILB_ = 2(*d*_HC_ + *d*_HG_). ^c^Not defined.

| Layer | $d$ [nm] ($\pm 0.1$) | $\rho$ [$\text{e}^-/\text{\AA}^3$] ($\pm 0.02$) |
| --- | --- | --- |
| OTS | 2.0 | 0.34 |
| MeD | 0.4 | 0.17 |
| HC | 1.8 | 0.30 |
| HG | 0.9 | 0.40 |
| W | 0.4 | 0.33<sup>a</sup> |
| ILB | 5.4<sup>b</sup> | (-)<sup>c</sup> |

In conclusion, the electron density profile reconstructed from the XRR data confirms that the samples have the intended architecture suited for measurements of pressure–distance curves by ellipsometry. The interstitial water layer thickness as determined by XRR at ambient humidity is *d*_W_ ≈ 0.4 nm, and the total organic thickness of the inverse bilayer is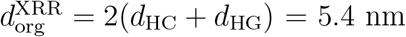. Both values serve for a comparison with the results of the ellipsometry measurements presented next.

### Pressure–distance curves: ellipsometry

We move on with the sample characterization by ellipsometry performed under dry conditions and in a wide range of hydration levels. The faster ellipsometry was chosen for this purpose instead of XRR because XRR measurements take too long to keep up with the changes in humidity, which occur on the time scales of minutes or less. For the analysis of the ellipsometric angles we assumed *n*_Si_ = 3.885–0.018i as the complex refractive index of silicon^37^ and *n*_oxi_ = 1.46 for the refractive index of SiO_2_, as determined earlier.^24^ The ellipsometric angles obtained for fully dehydrated samples (*h*_rel_ ≲ 3%, achieved by streaming with dry N_2_) were modeled by accounting for the oxide layer properties determined before, to obtain the thickness *D*_org_ of the dry organic layer jointly formed by OTS and the two lipid monolayers. In this procedure, we assumed a refractive index of *n*_org_ = 1.5, in line with earlier reports on organic materials.^38,39^ This analysis resulted in *D*_OTS_ = *D*_org_(OTS) = 1.4 ± 0.1 nm and *D*_tot_ = *D*_org_(OTS+ILB) = 7.2 ± 0.9 nm, from which the ILB thickness can be deduced as *D*_ILB_ = *D*_tot_ − *D*_OTS_ = 5.8 ± 0.8 nm. The values are summarized in Table 2, which also contains the ellipsometric angles Ψ that are most sensitive to the organic layer thickness under the given conditions. The thicknesses of OTS and of the ILB, as obtained by ellipsometry, are in reasonable agreement with the ones obtained by XRR (*d*_OTS_ ≈ 2 nm, *d*_ILB_ ≈ 5 nm, see Table 1), where the uncertainty mainly arises from the uncertainty in the choice of *n*_org_. It should, however, be noted that no such uncertainty applies to the water layer dealt with in the following, because the refractive index of water is precisely known and for visible light is virtually independent of its supramolecular organization. ^40–42^

**Table 2:** Ellipsometric angle Ψ and associated thickness values *D*_OXI_ and *D*_org_ from which the thicknesses of OTS and of the ILB, *D*_OTS_ and *D*_ILB_, respectively, are deduced as described in the main text. Uncertainties in the thicknesses are estimated as ± 0.1 nm.

| Bare oxide: | $\Psi(\text{OXI}) = 41.2^\circ$ | $D_{\text{OXI}} = 99.8$ nm | | | |
| --- | --- | --- | --- | --- | --- |
| ILB sample | $\Psi(\text{OXI} + \text{OTS}) [^\circ]$ | $D_{\text{OTS}}$ [nm] | $\Psi(\text{OXI} + \text{OTS} + \text{ILB}) [^\circ]$ | $D_{\text{tot}}$ [nm] | $D_{\text{ILB}}$ [nm] |
| DSPC-DSPC | 42.1 | 1.4 | 45.0 | 6.3 | 4.9 |
| DSPC-DGDG | 42.1 | 1.3 | 45.5 | 6.5 | 5.2 |
| DGDG-DSPC | 42.2 | 1.5 | 46.1 | 7.9 | 6.4 |
| DGDG-DGDG | 42.3 | 1.5 | 46.1 | 8.0 | 6.5 |

In the last step, the ellipsometric angles, obtained at controlled humidity *h*_rel_, were modeled while accounting for the known optical parameters of the oxide and of the dry organic layers. The humidity-dependent water layer thickness *D*_w_(*h*_rel_) was then determined assuming *n*_w_ = 1.33 as the refractive index of water. Note that *D*_w_ should be understood as an equivalent thickness in terms of the *z*-integrated water density profile rather than as a distinct layer of pure water.^24^ The position of the water layer with respect to the organic layer in the model has negligible influence on the results because in the thin-film limit ellipsometry is merely sensitive to the overall optical path.^24^ A dedicated humidity control setup (see Methods section) enabled ellipsometry measurements in a wide range of controlled humidities, 4% ≲ *h*_rel_ ≲ 97%, and allowed us to probe an interaction pressure range of 4.0 · 10^6^ Pa ≲ Π ≲ 4.0 · 10^8^ Pa. Increasing the humidity above 98% resulted in condensation on the outer sample surface, which not only prevents any meaningful analysis of the ellipsometric angles but may also affect the integrity of the ILB architecture.

Fig. 3 shows the obtained pressure–distance curves Π(*D*_w_) of various interacting lipid membrane surfaces in a semi-logarithmic representation. The dehydrating interaction pressure Π was calculated from the measured relative humidity according to Eq. 2. At Π = 3.0 · 10^8^ Pa, the layers were assumed to be maximally dehydrated, so that this pressure was used for the definition of *D*_w_ = 0. It should be noted, however, that even under these extremely dry conditions around one water molecules per lipid may persist. ^43^ The definition of *D*_w_ = 0 was chosen because Π = 3.0 · 10^8^ Pa is around the maximal pressure reproducibly reached in the experiments. As shown in the Supporting Information S3, no water uptake is observed when the bare OTS-functionalized surfaces are exposed to varying humidities, confirming that the water uptake by the ILBs must be fully attributed to the interaction of the hydrophilic surfaces of the lipid layers, as intended.

**Figure 3:**
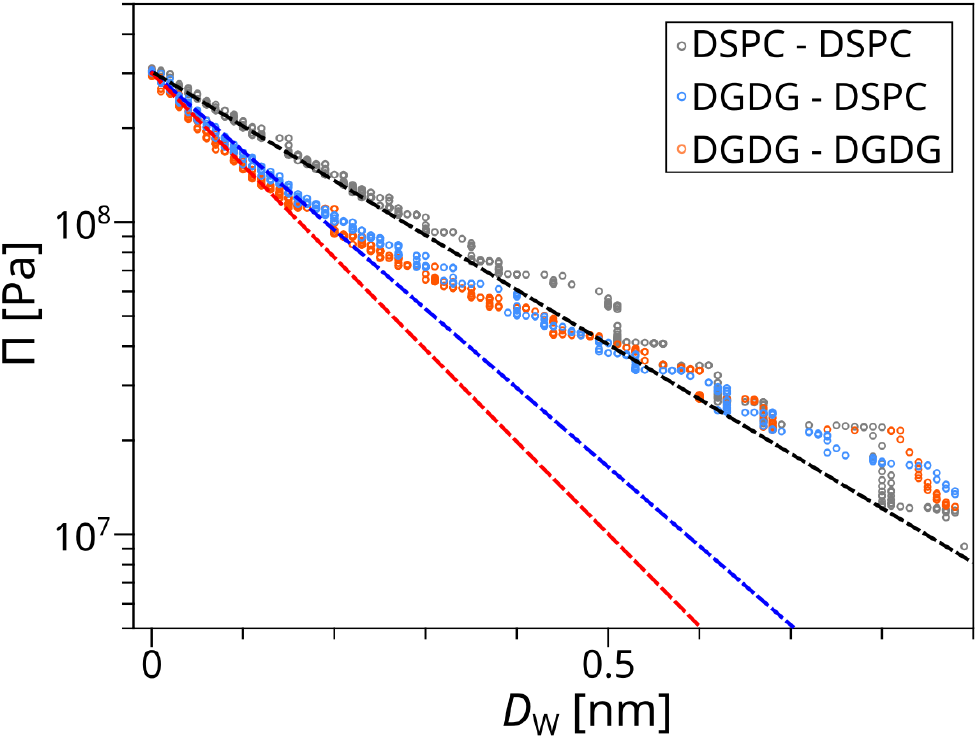
Pressure–distance curves obtained with symmetrical ILBs of either two PC lipid monolayers or two DGDG monolayers and with an asymmetrical ILB of one DGDG monolayer at the bottom and one PC lipid monolayer on top. Data from an asymmetrical ILB with the reverse sequence are shown in the Supporting Information (Fig. S4). Dashed lines indicate fits to the initial exponential decay of the curves.

In the case of two interacting identical PC lipid layers, realized here with an ILB entirely composed of DSPC, the pressure–distance curve follows an approximately exponential decay, in line with the literature data on PC lipid membranes that are mostly based on experiments with multilayers. ^3,10,12^ At ambient humidity (typically *h*_rel_ ≈ 30% - 50%) a water layer thickness of ≈ 0.2-0.4 nm is expected according to the pressure–distance curve. This thickness is in good agreement with *d*_W_ ≈ 0.4 nm, as obtained by XRR, see Table 1. An exponential fit to the pressure–distance data of the form

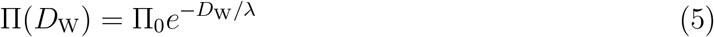

yields the characteristic decay length *λ*. Here, a linear fit was made to the natural logarithm of the pressure data in order to give equal weights to all data points. In Fig. 3, this fit is shown as a straight black line. The value obtained for two interacting PC lipid surfaces, *λ* = 0.26 nm, is in satisfactory agreement with the consensus value for chain-ordered PC lipid bilayers established by Kowalik et al., ^16^ *λ* = 0.21 nm. The decay lengths obtained for all ILBs are summarized in Table 3. The values of Π_0_ are not reported because Π_0_ = 3.0 · 10^8^ Pa (by construction) and therefore should not be directly compared with Π_0_ reported in multilayer-based studies.^12,18^

**Table 3:** Decay lengths of the pressure–distance curves of symmetrical and asymmetrical ILBs as obtained in exponential fits and comparison with literature data. ^a^Limited comparability because of different phase.

| Sample | DSPC-DSPC | DGDG-DGDG | DGDG-DSPC |
| --- | --- | --- | --- |
| $\lambda$ [nm] ($\pm 0.01$) | 0.26 | 0.15 | 0.17 |
| $\lambda$ literature [nm] | 0.21 (ref. <sup>16</sup>) | 0.12<sup>a</sup> (ref. <sup>17</sup>) | - |

We move on with the pressure–distance curve of an ILB composed of two interacting identical DGDG layers. It is seen in Fig. 3 that Π(*D*_w_) initially drops more rapidly with increasing water layer thickness than for the interacting PC lipid surfaces, confirming that the chemical details govern the pressure–distance curve at very short separation. For larger separations the curve becomes again more similar to that of the two interacting PC lipid layers, suggesting that the repulsion mechanisms are less chemistry-specific at larger separations. An exponential fit in the range of low *D*_W_ yields a decay length of only *λ* = 0.15 nm, which is in good agreement with the decay length of 0.12 nm reported previously for interacting DGDG membranes.^3,17^ The two values should, however, be compared with caution, because in those earlier studies with membrane multilayers the membranes were in the fluid state characterized by a larger area per molecule because of a different chain chemistry featuring numerous double bonds. Nevertheless, our present results confirm the clear trend that the decay length is much shorter for interacting glycolipid membrane surfaces than for interacting PC lipid membrane surfaces.

Finally, we turn to the pressure–distance curve of an asymmetrical ILB composed of one DSPC layer interacting with a DGDG layer, recalling that such a scenario involving two dissimilar lipid surfaces cannot be accessed with conventional approaches for the reasons specified in the introduction. As seen in Fig. 3, the associated Π(*D*_w_) decays almost as rapidly as that of the symmetrical ILB of two DGDG layer. In fact, the decay length obtained in the exponential fit, *λ* = 0.17 nm, is only slightly larger. As shown in the Supporting Information, a consistent pressure–distance curve is obtained when inverting the sequence of deposition, i.e., first transferring a DSPC monolayer and then transferring a DGDG monolayer on top of it. The minor differences between the curves may be attributed to possible trace amounts of lipids from the first monolayer remaining on the water surface, so that the two scenarios may not be perfectly equivalent.

The observation of such a small decay length for the asymmetrical ILBs of DSPC and DGDG appears puzzling at first, because a more intermediate behavior (i.e., half way between the pressure–distance curves of the two symmetrical ILBs) may be expected naively. In the following, we will however provide an explanation based on the current knowledge of the physical mechanisms of the hydration repulsion.

Phospholipids like the PC lipids studied here have zwitterionic headgroups featuring a single large electric dipole^44^ with orientational degrees of freedom. With regard to the hydration repulsion between phospholipid membrane surfaces in water, apart from the release of the most strongly bound water molecules,^45^ two mechanisms have been proposed, both of which are ultimately related to this dipolar headgroup architecture. The first one is a consequence of water structuring effects via the strong orientational polarization of the water layers interacting with the array of oriented headgroup dipoles displayed at the membrane surfaces. The repulsion then results from an unfavorable overlap of the opposing order parameter profiles, which increases with decreasing surface separation. ^15^ This picture has later been refined^46–49^ and recently scrutinized on a quantitative level through direct comparison with observables extracted from MD simulations. ^26^ The second mechanism has to do with the configurational entropy of the lipids and its unfavorable decrease upon surface approach. ^7,50^ For PC lipids there is a pronounced entropy loss due to the strong directional coupling of the electric dipoles in the opposing membrane surfaces. ^19^

It has, however, been shown that both of these repulsion mechanisms are essentially inoperative for lipids with saccharide headgroups,^17^ like the DGDG studied here. The reason is that saccharide headgroups do not have a large single electric dipole but instead feature many small electric dipoles in the form of -OH groups that are oriented in all possible directions. As a consequence, no significant orientational polarization of the interfacial water occurs that would lead to water-structural hydration repulsion. Moreover, it was found in the same study that the orientation of DGDG headgroups virtually does not respond to variations of the membrane proximity,^17^ so that no entropic repulsion of that sort would arise either. In essence, the -OH groups displayed by the DGDG surfaces behave similarly to those of water. For the asymmetrical case of PC lipid layers interacting with DGDG lipid layers one may thus expect that there is little interference between the orientationally polarized water layers adjacent to the PC lipid surface and the hydrated saccharide layer belonging to the DGDG surface. Likewise, no directional coupling is expected to occur between the large PC dipoles of the one surface and the saccharide headgroups of the other surface. These conclusions rationalize why the comparatively long-range hydration repulsion is only observed in the symmetrical case of two PC lipid surfaces, whereas the underlying repulsion mechanisms become insignificant as soon as one of the surfaces is replaced with a glycolipid layer.

At this point we would like to emphasize again the excellent comparability of the experimental data obtained here using ILBs with data obtained using atomistic MD simulations, because of the consistent definition of the water layer thickness and because of the overlapping ranges of accessible Π and *D*_W_ values.^16,17,19,26^ The present results therefore strongly motivate MD simulations of the hydration repulsion between dissimilar lipid membrane surfaces. Such future simulations, which would go beyond the scope of the present experimental study, clearly have the potential to elucidate the atomistic details of the underlying interaction mechanisms, once validated with experimental pressure–distance curves.

## Conclusions

The inverse lipid bilayer (ILB) architecture determined by XRR in combination with ellipsometry under controlled humidity conditions offers a versatile measurement platform for the determination of pressure–distance curves between identical and dissimilar hydrophilic surfaces of amphiphilic layers. The pressure–distance curves are quantitative and with regard to *D*_W_ and the measurements can be performed routinely in a laboratory without X-ray or neutron source and with extremely small sample amounts, which may be important for expensive materials.

In the present work, we were able to reproduce the known pressure–distance curves of identical phospholipid and glycolipid surfaces at short separations and provided first sets of data on the interaction between two dissimilar membrane surfaces, with phospholipids on one side and glycolipids on the other side. In the future, this experimental work should be followed by MD simulation studies predicting the pressure–distance curves of such asymmetrical interaction scenarios, as was done successfully for symmetrical interaction scenarios in the past.^16,17^ Such simulations will likely yield important insights into the physical mechanisms responsible for the repulsive short-range interaction between hydrophilic surfaces of various kinds under water.

## Materials and methods

### Chemicals, lipids, and solid substrates

Chloroform and methanol (purity ≥ 99.9%, [Warning: chloroform and methanol are toxic on incorporation, inhalation, skin, and eye contact; use under fume hood and/or with suitable personal protection]) were purchased from Merck (Darmstadt, Germany) and used without further purification. OTS was purchased from abcr GmbH (Karlsruhe, Germany). Double-deionized ultrapure water (resistivity: 18.2 MΩ·cm) was obtained from a water purification station (Purelab classic, Elga, Celle, Germany). All lipids used were purchased from Avanti Polar Lipids (Alabaster, Alabama, United States) or its distributor Merck. Silicon chips were cut from larger wafers. The silicon wafers (150 mm diameter, 625 *µ*m thickness) of which the polished surface was covered with layer of thermal silicon oxide of 100 nm nominal thickness were purchased from SIEGERT Wafer GmbH (Aachen, Germany) and cut into rectangular pieces of ≈ 30 mm × 20 mm (termed chips throughout this manuscript).

### Sample preparation

Glassware and sample vials were washed with chloroform before use. Lipid solutions were prepared by dissolving lipid powder at a concentration of 1 mg/mL in chloroform or a mixture of chloroform and methanol (2:1 by volume), for phospholipid or glycolipids, respectively. The silicon chips used in the experiments were first thoroughly cleaned and the hydrophobically functionalized with a layer of OTS to obtain a hydrophobic surface. The whole procedure is detailed in the Supporting Information. The hydrophobized chips were rinsed with chloroform and ethanol before use.

Inverse lipid bilayers (ILBs) were prepared at room temperature by the sequential deposition of two individual lipid monolayers from the air/water interface onto the surfaces of the chips with the Langmuir-Blodgett (LB) technique. A teflon Langmuir trough by Riegler & Kirstein (Potsdam, Germany) was used for this purpose. Starting with the bare hydrophobized chip outside the aqueous medium, both transfers were performed with LB, by moving the chip first in, then out of the medium, at a constant speed of 6 mm/min. Before the first transfer, a lipid monolayer at the air/water interface was prepared by spreading a suitable amount of lipid solution, allowing 10 min for complete solvent evaporation, and compressing the monolayer to the desired lateral pressure of *π* = 40 mN/m, which was kept constant during the transfers through barrier movements. The lateral pressure was monitored by a Wilhelmy-plate. Asymmetrical ILBs were prepared by replacing the initial lipid monolayer from the air/water interface and spreading a new one with a different lipid composition between the first and second transfers.

### X-ray reflectometry

XRR measurements were performed using a D8 Advance reflectometer (Bruker AXS, Karlsruhe, Germany), as described in reference,^33^ from which the following paragraph is largely reproduced. Reflectivity curves were measured in the *θ*–2*θ* geometry, where *θ* is the incident angle. A conventional X-ray tube with a Cu anode (Cu K*α*, wavelength *λ* = 1.54 Å) was used to generate an X-ray beam with a line focus. The primary beam was monochromatized by a Göbel mirror (W/Si multilayer mirror) and collimated in the *z*-direction through two narrow slits of 0.1 mm each with a switchable absorber (calibrated Cu attenuator) in between them. Additionally a Soller-slit (1^*°*^ divergence) collimates the primary beam in the *y*-direction. After scattering, on the secondary arm, the same slit collimation as on the primary side was employed. The intensity was recorded with an O-D scintillation counter (Bruker AXS). Data were corrected using the known attenuation factors and counting times. Finally, the angular reflectivity scans were transformed to reflectivity curves as a function of the perpendicular scattering vector component, *q*_*z*_ = 4*π/λ* sin *θ*, and corrected for the footprint effect. For analysis, the experimental data were compared with theoretically modeled XRR curves based on a slab-model representation of the electron density profile at the interface between air and the silicon bulk material (see Results section). These profiles were discretized into 1-Å-thin sub-slabs of constant electron density, and the corresponding *q*_*z*_-dependent reflectivities, *R*(*q*_*z*_), were then calculated from the Fresnel reflection laws at each slab-slab interface using the iterative recipe of Parratt.^51^ Finally, all model parameters (electron densities, layer thicknesses, and roughness) were varied until the best agreement with the experimental data was reached via *χ*^2^ minimization. Estimates of the statistical parameter errors, e.g. corresponding to the 95% (two-Sigma) confidence interval, are valid only within the framework of a “perfect model”, characterized by a reduced *χ*^2^ close to unity^52^ and typically greatly underestimate the real uncertainty. In view of significant additional contributions due to systematic errors, much larger error estimates are therefore provided instead in the tables next to the parameter values. They approximately reflect the variation of the obtained parameters throughout the evolution and refinement of the above-described model description, i.e., they reflect the robustness of the parameters with respect to the model, and we therefore consider them more meaningful.^24^

### Ellipsometry

As described in an earlier publication,^24^ from which the following paragraph is partially reproduced, ellipsometry enables the characterization of interfacial layers in terms of refractive indices and thicknesses. The method is based on the change in the polarization state of light upon reflection from the surface. For a given refractive index *n*, the change depends on the layer thickness and is quantified in terms of the phase difference Δ and the amplitude ratio Ψ encoded in the ratio between the complex reflection coefficients *R*_s_ and *R*_p_ for s and p polarizations, respectively, ^53^

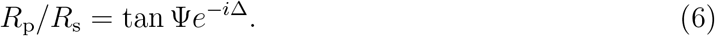

Ellipsometry measurements were carried out before and after the ILB depositions. The chips used as substrate have a rather thick (≈ 1000 Å) layer of thermally annealed oxide on their surface. The measurements were performed at an incident angle of 70^*°*^ with a Multiskop ellipsometer (Optrel, Berlin, Germany) working with a wavelength *λ*_elli_ = 632.8 nm.

### Humidity control chamber

To observe the water uptake into the ILBs in response to variations in the relative humidity (i.e., in the chemical potential of water), the ellipsometry measurements were conducted in a humidity control chamber (HCC, see Fig. 4). The inflow was fed by the in-house dry nitrogen (N_2_) supply and split into two branches that reconnect before reaching the HCC. One branch is saturated with moisture through sequential bubbling through two water bottles while the other branch remains dry. Adjusting the ratio of dry and humidified N_2_ allows for continuous variations in the resulting humidity. In order to reach the highest humidities (up to 97%) the flux of dry N_2_ was stopped completely, and the water in the bottles was gently heated above room temperature. The relative humidity in the chamber was measured with a HYTELOG USB sensor (B+B Thermo-Technik, Donaueschingen, Germany) with a measurement range 0 − 100%, a resolution of 0.01% and an accuracy of ±2%.

**Figure 4:**
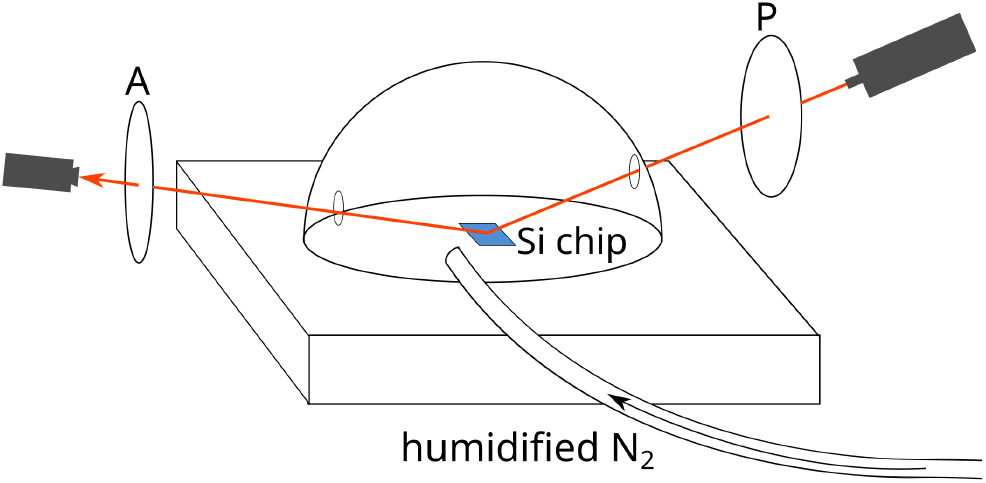
Schematic illustration of the humidity control chamber used for ellipsometry in this study. The laser path is illustrated with red lines through the polarizer (P) and the analyzer (A).

### A-posteriori synchronization of humidity and ellipsometry data

Ellipsometer and humidity sensor record their measurements independently, so that the data had to be synchronized a-posteriori. This was achieved by linear interpolation of the humidity measurement values recorded at a frequency 0.5 Hz at the time points of the ellipsometric measurements, which were recorded at a frequency of 0.1 Hz. This procedure allowed us eliminating the time as parameter and reconstructing the sought pressure–distance curve Π(*D*_W_).

## Supporting information

Functionalization with OTS; Full Set of XRR Parameters; Second XRR measurement; PD-curve of a blank OTS surface; Comparison between asymmetrical ILBs with different deposition sequence of DSPC and DGDG.

## Acknowledgement

We thank Harald Hartig and Ricardo Gioia-Alvarez for help with the experimental setup for ellipsometry under controlled humidity.

## References

(1) Alberts, B.; Bray, D.; Lewis, J.; Raff, M.; Roberts, K.; Watson, J. D.; others Molecular biology of the cell; Garland New York, 1994; Vol. 3.

(2) Lipowsky, R.; Sackmann, E. Structure and dynamics of membranes: I. from cells to vesicles/II. generic and specific interactions; Elsevier, 1995.

(3) Demé, B.; Cataye, C.; Block, M. A.; Maréchal, E.; Jouhet, J. Contribution of galactoglycerolipids to the 3-dimensional architecture of thylakoids. The FASEB Journal 2014, 28, 3373–3383.

(4) Israelachvili, J. N. Intermolecular and surface forces; Academic press, 2015.

(5) Helfrich, W. Steric interaction of fluid membranes in multilayer systems. Zeitschrift für Naturforschung A 1978, 33, 305–315.

(6) Langmuir, I. The role of attractive and repulsive forces in the formation of tactoids, thixotropic gels, protein crystals and coacervates. The Journal of Chemical Physics 1938, 6, 873–896.

(7) Israelachvili, J. N.; Wennerstroem, H. Entropic forces between amphiphilic surfaces in liquids. The Journal of Physical Chemistry 1992, 96, 520–531.

(8) Klitzing, R. v. Effect of interface modification on forces in foam films and wetting films. Advances in colloid and interface science 2005, 114, 253–266.

(9) Lipowsky, R. The conformation of membranes. Nature 1991, 349, 475–481.

(10) LeNeveu, D.; Rand, R. P.; Parsegian, V. A. Measurement of forces between lecithin bilayers. Nature 1976, 259, 601–603.

(11) Marra, J.; Israelachvili, J. Direct measurements of forces between phosphatidylcholine and phosphatidylethanolamine bilayers in aqueous electrolyte solutions. Biochemistry 1985, 24, 4608–4618.

(12) Parsegian, V. A.; Fuller, N.; Rand, R. P. Measured work of deformation and repulsion of lecithin bilayers. Proceedings of the National Academy of Sciences 1979, 76, 2750–2754.

(13) Leontidis, E.; Aroti, A.; Belloni, L.; Dubois, M.; Zemb, T. Effects of monovalent anions of the Hofmeister series on DPPC lipid Bilayers part II: Modeling the perpendicular and lateral equation-of-state. Biophysical journal 2007, 93, 1591–1607.

(14) Schneck, E.; Demé, B.; Gege, C.; Tanaka, M. Membrane Adhesion via homophilic saccharide-saccharide interactions investigated by neutron scattering. Biophysical Journal 2011, 100, 2151–2159.

(15) Marčelja, S.; Radić, N. Repulsion of interfaces due to boundary water. Chemical Physics Letters 1976, 42, 129–130.

(16) Kowalik, B.; Schlaich, A.; Kanduc, M.; Schneck, E.; Netz, R. R. Hydration repulsion difference between ordered and disordered membranes due to cancellation of membrane– membrane and water-mediated interactions. The Journal of Physical Chemistry Letters 2017, 8, 2869–2874.

(17) Kanduč, M.; Schlaich, A.; de Vries, A. H.; Jouhet, J.; Demé, B.; Netz, R. R.; Schneck, E. Tight cohesion between glycolipid membranes results from balanced water–headgroup interactions. Nat. Comm. 2017, 8.

(18) Lis, L.; McAlister, D.; Fuller, N.; Rand, R.; Parsegian, V. Interactions between neutral phospholipid bilayer membranes. Biophysical Journal 1982, 37, 657–665.

(19) Schneck, E.; Sedlmeier, F.; Netz, R. R. Hydration repulsion between biomembranes results from an interplay of dehydration and depolarization. Proceedings of the National Academy of Sciences 2012, 109, 14405–14409.

(20) Marra, J. Controlled deposition of lipid monolayers and bilayers onto mica and direct force measurements between galactolipid bilayers in aqueous solutions. Journal of colloid and interface science 1985, 107, 446–458.

(21) Yu, Z.; Calvert, T.; Leckband, D. Molecular forces between membranes displaying neutral glycosphingolipids: evidence for carbohydrate attraction. Biochemistry 1998, 37, 1540–1550.

(22) Israelachvili, J.; Min, Y.; Akbulut, M.; Aligl, A.; Carver, G.; Greene, W.; Kristiansen, K.; Meyer, E.; Pesika, N.; Rosenberg, K.; Zeng, H. Recent advances in the surface forcesapparatus (SFA) technique. Reports on Progress in Physics 2010, 73.

(23) Schneider, M. F.; Mathe, G.; Tanaka, M.; Gege, C.; Schmidt, R. R. Thermodynamic properties and swelling behavior of glycolipid monolayers at interfaces. The Journal of Physical Chemistry B 2001, 105, 5178–5185.

(24) Rodriguez-Loureiro, I.; Scoppola, E.; Bertinetti, L.; Barbetta, A.; Fragneto, G.; Schneck, E. Neutron reflectometry yields distance-dependent structures of nanometric polymer brushes interacting across water. Soft Matter 2017, 13, 5767–5777.

(25) Schneck, E.; Rodriguez-Loureiro, I.; Bertinetti, L.; Marin, E.; Novikov, D.; Konovalov, O.; Gochev, G. Element-specific density profiles in interacting biomembrane models. Journal of Physics D: Applied Physics 2017, 50, 104001.

(26) Schlaich, A.; Daldrop, J. O.; Kowalik, B.; Kanduč, M.; Schneck, E.; Netz, R. R. Water structuring induces nonuniversal hydration repulsion between polar surfaces: quantitative comparison between molecular simulations, theory, and experiments. Langmuir 2024, 40, 7896–7906.

(27) Stefaniu, C.; Latza, V. M.; Gutowski, O.; Fontaine, P.; Brezesinski, G.; Schneck, E. Headgroup-Ordered Monolayers of Uncharged Glycolipids Exhibit Selective Interactions with Ions. The Journal of Physical Chemistry Letters 2019, 10, 1684–1690.

(28) Mukhina, T.; Brezesinski, G.; Shen, C.; Schneck, E. Phase behavior and miscibility in lipid monolayers containing glycolipids. Journal of Colloid and Interface Science 2022, 615, 786–796.

(29) Nagata, K.; Ogure, A.; Hirosawa, I.; Suwa, T.; Teramoto, A.; Hattori, T.; Ohmi, T. Structural Analyses of Thin SiO2 Films Formed by Thermal Oxidation of Atomically Flat Si Surface by Using Synchrotron Radiation X-Ray Characterization. ECS Journal of Solid State Science and Technology, 2015, N96–N98.

(30) Pomerantz, M.; Segmüller, A.; Netzer, L.; Sagiv, J. Coverage of Si substrates by self-assembling monolayers and multilayers as measured by IR, wettability and X-ray diffraction. Thin Solid Films 1985, 132, 153–162.

(31) Mezger, M.; Reichert, H.; Schöder, S.; Okasinski, J.; Schröder, H.; Dosch, H.; Palms, D.; Ralston, J.; Honkimäki, V. High-resolution in situ x-ray study of the hydrophobic gap at the water–octadecyl-trichlorosilane interface. Proceedings of the National Academy of Sciences 2006, 103, 18401–18404.

(32) Schneck, E.; Berts, I.; Halperin, A.; Daillant, J.; Fragneto, G. Neutron reflectometry from poly (ethylene-glycol) brushes binding anti-PEG antibodies: Evidence of ternary adsorption. Biomaterials 2015, 46, 95–104.

(33) Pusterla, J.; Scoppola, E.; Appel, C.; Mukhina, T.; Shen, C.; Brezesinski, G.; Schneck, E. Characterization of lipid bilayers adsorbed to functionalized air/water interfaces. Nanoscale 2022, 14, 15048–15059.

(34) Petrache, H. I.; Tristram-Nagle, S.; Nagle, J. F. Fluid phase structure of EPC and DMPC bilayers. Chemistry and Physics of Lipids 1998, 95, 83–94.

(35) Bange, L.; Mukhina, T.; Fragneto, G.; Rondelli, V.; Schneck, E. Influence of adhesion-promoting glycolipids on the structure and stability of solid-supported lipid double-bilayers. Soft Matter 2024, 20, 2113–2125.

(36) Kjaer, K.; Als-Nielsen, J.; Helm, C. A.; Tippman-Krayer, P.; Moehwald, H. Synchrotron x-ray diffraction and reflection studies of arachidic acid monolayers at the air-water interface. The Journal of Physical Chemistry 1989, 93, 3200–3206.

(37) Adachi, S. Model dielectric constants of Si and Ge. Physical Review B 1988, 38, 12966.

(38) Wong, J. E.; Rehfeldt, F.; Hänni, P.; Tanaka, M.; Klitzing, R. v. Swelling behavior of polyelectrolyte multilayers in saturated water vapor. Macromolecules 2004, 37, 7285– 7289.

(39) Ruths, J.; Essler, F.; Decher, G.; Riegler, H. Polyelectrolytes I: polyanion/polycation multilayers at the air/monolayer/water interface as elements for quantitative polymer adsorption studies and preparation of hetero-superlattices on solid surfaces. Langmuir 2000, 16, 8871–8878.

(40) Schiebener, P.; Straub, J.; Levelt Sengers, J.; Gallagher, J. Refractive index of water and steam as function of wavelength, temperature and density. Journal of physical and chemical reference data 1990, 19, 677–717.

(41) Le, T.; Morita, A.; Tanaka, T. Refractive index of nanoconfined water reveals its anomalous physical properties. Nanoscale horizons 2020, 5, 1016–1024.

(42) McKee, C. T.; Ducker, W. A. Refractive index of thin, aqueous films between hydrophobic surfaces studied using evanescent wave atomic force microscopy. Langmuir 2005, 21, 12153–12159.

(43) Markova, N.; Sparr, E.; Wadsö, L.; Wennerström, H. A calorimetric study of phospholipid hydration. Simultaneous monitoring of enthalpy and free energy. The Journal of Physical Chemistry B 2000, 104, 8053–8060.

(44) Wohlert, J.; Edholm, O. The Range and Shielding of Dipole-Dipole Interactions in Phospholipid Bilayers. Biophysical Journal 2004, 87, 2433–2445.

(45) Israelachvili, J.; Wennerström, H. Role of hydration and water structure in biological and colloidal interactions. Nature 1996, 379, 219–225.

(46) Radić, N.; Marčelja, S. Solvent contribution to the debye screening length. Chemical Physics Letters 1978, 55, 377–379.

(47) Cevc, G.; Podgornik, R.; Zeks, B. The free energy, enthalpy and entropy of hydration of phospholipid bilayer membranes and their difference on the interfacial separation. Chemical Physics Letters 1982, 91, 193–196.

(48) Ruckenstein, E.; Schiby, D. On the origin of repulsive hydration forces between two mica plates. Chemical Physics Letters 1983, 95, 439–443.

(49) Attard, P.; Batchelor, M. T. A mechanism for the hydration force demonstrated in a model system. Chemical physics letters 1988, 149, 206–211.

(50) Israelachvili, J. N.; Wennerstroem, H. Hydration or steric forces between amphiphilic surfaces? Langmuir 1990, 6, 873–876.

(51) Parratt, L. G. Surface studies of solids by total reflection of X-rays. Physical Review 1954, 95, 359.

(52) Bevington, P. R.; Robinson, D. K.; Blair, J. M.; Mallinckrodt, A. J.; McKay, S. Data reduction and error analysis for the physical sciences. Computers in Physics 1993, 7, 415–416.

(53) Azzam, R. M.; Bashara, N. M.; Ballard, S. S. Ellipsometry and polarized light. Physics Today 1978, 31, 72.

