## supporting information for "Pressure–distance curves of identical and dissimilar lipid membrane surfaces in water"

#### Functionalization with OTS

To turn the hydrophilic $SiO_2$ surface of the silicon chips hydrophobic, the chips were covalently functionalized with a layer of octadecyltrichlorosilane (OTS, see fig. S1). For this purpose, the silicon chips were first thoroughly cleaned by rinsing them with a cascade of solvents including chloroform, acetone, ethanol, and water for 10 min per solvent. Subsequently, the chips were rinsed again with ethanol to remove remaining traces of the water and to facilitate drying with a stream of dry nitrogen. The dry chips were then treated for another 15 min in an OV/ozone cleaner (Bioforce Nanosciences, Ames, Iowa, United States). They were then placed in a 1 mM solution of OTS in anhydrous toluene for 60 min to allow the covalent attachment of the OTS to the silicon surface to take place. Finally toluene residues were removed by two rinsing and drying cycles with ethanol. The success of the hydrophobization procedure was verified by observing the high contact angle of a water droplet placed on the surface. The hydrophobic functionalization with OTS was stable over a period of several months, allowing the chips to be reused several times. The integrity of the OTS layer was verified before every measurement by observing the water contact angle and by ellipsometric measurements of the OTS layer thickness.

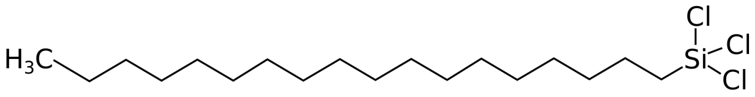


Figure S1: Chemical structure of octadecyltrichlorosilane (OTS)

#### Full Set of XRR Parameters

Table S1: Structural parameters of a DSPC ILB on OTS at ambient humidity as deduced from the XRR experiments. [a] $d_{\mathrm{IBL}} = 2(d_{\mathrm{HC}} + d_{\mathrm{HG}})$.

| **layer** | $d$ [nm] | $\rho$ [e$^-$/Å$^3$] | transition | $\sigma$ [nm] |
|---|---|---|---|---|
| $SiO_2$ | - | 0.73 | $SiO_2$ - OTS | 0.4 |
| OTS | 2.0 | 0.34 | OTS - MD | 0.2 |
| MD | 0.4 | 0.17 | MD - HC | 0.6 |
| HC | 1.8 | 0.30 | HC - HG | 0.1 |
| HG | 0.9 | 0.40 | HG - W | 0.6 |
| W | 0.4 | 0.33 | HC - AIR | 0.5 |
| IBL | 5.4[a] | - | - | |

#### Reflectivity data of the second ILB sample prepared for XRR measurements

Fig. S2 shows the reflectivity data of the second DSPC ILB sample prepared with the same protocol. It is virtually identical to the reflectivity data from the first sample (shown in the main text). The solid line is the simulated reflectivity curve corresponding to the parameters in Table S1, which also reproduces the reflectivity data from the first sample. This result confirms that the sample preparation is highly reproducible.

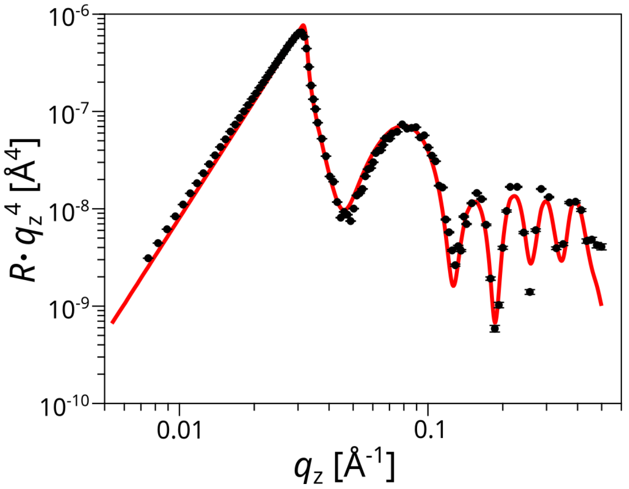


Figure S2: Reflectivity data of the second DSPC ILB sample prepared with the same protocol. Solid line: Simulated reflectivity curve corresponding to the parameters obtained with the first sample (Table S1).

#### Apparent pressure–distance curve of a bare chip

Fig. S3 shows the apparent pressure-distance curve measured with a bare OTS-functionalized chip without ILB. It is seen that the hydrophobic surface does not adsorb any significant water layer when the humidity increases.

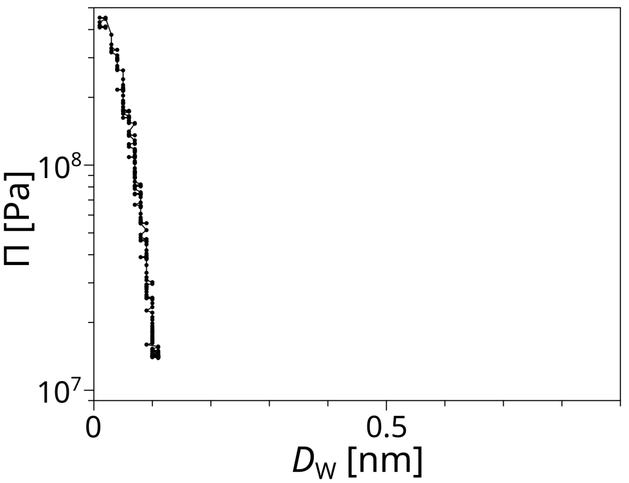


Figure S3: Apparent pressure–distance curve of a bare OTS-functionalized chip without ILB.

### Comparison of pressure–distance curves from asymmetrical ILBs with reversed monolayer order

Fig. S4 shows the pressure–distance curves of asymmetric ILBs. In the sample denoted with "DGDG-DSPC" in the figure legend, a monolayer of the glycolipid DGDG was deposited first and a DSPC monolayer was deposited on top of it afterwards. For "DSPC-DGDG" that sequence is reversed with DSPC being deposited first. The overall trend of the two pressure–distance curves is very similar, as reflected also in the similar decay lengths displayed in Table S2.

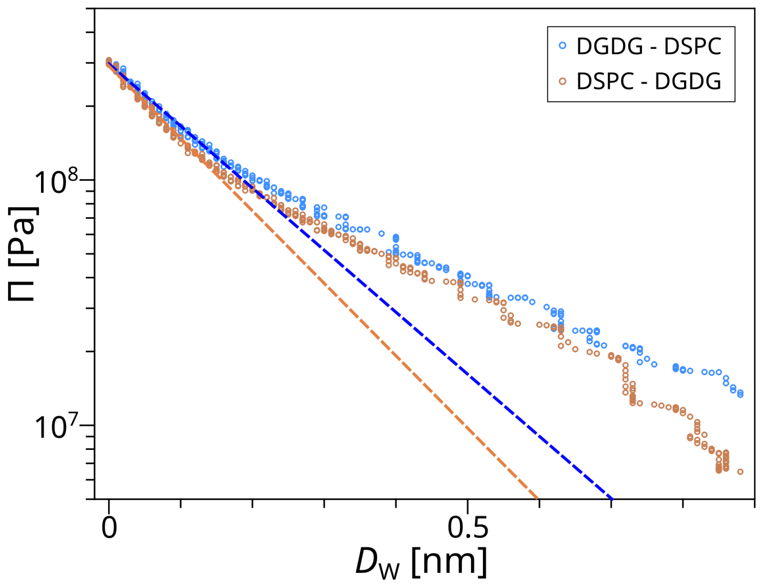


Figure S4: Pressure–distance curves obtained with two different asymmetrical ILBs of one PC lipid monolayer and one DGDG monolayer. They differ in the sequence of the deposited monolayers, as indicated in the legend. The dashed lines indicate fits to the initial exponential decays of the data points.

Table S2: Decay lengths of the pressure–distance curves of two different asymmetrical ILBs as obtained in exponential fits.

| **Sample** | DGDG-DSPC | DSPC-DGDG |
|---|---|---|
| $\lambda$ [nm] ($\pm$ 0.01) | 0.17 | 0.15 |
